# Observed but Never Experienced – Vicarious Learning of Fear Under Ecological Conditions

**DOI:** 10.1101/2020.01.29.924720

**Authors:** Michał Szczepanik, Anna M. Kaźmierowska, Jarosław M. Michałowski, Marek Wypych, Andreas Olsson, Ewelina Knapska

**Affiliations:** Laboratory of Brain Imaging, Nencki Institute of Experimental Biology of Polish Academy of Sciences, 3 Pasteur Str., 02-093 Warsaw, Poland; Laboratory of Emotions Neurobiology, BRAINCITY - Centre of Excellence for Neural Plasticity and Brain Disorders, Nencki Institute of Experimental Biology of Polish Academy of Sciences, 3 Pasteur Str., 02-093 Warsaw, Poland; SWPS University of Social Sciences and Humanities, 10 Kutrzeby Str., 61-719 Poznań, Poland; Department of Clinical Neuroscience, Division of Psychology, Karolinska Institutet, Stockholm, Sweden

**Keywords:** observational fear learning, ecological validity, fear-potentiated startle, contingency learning, fear conditioning

## Abstract

Learning to avoid threats often occurs by observing the behavior of others. Most previous research on observational fear learning in humans has used pre-recorded stimuli as social cues. Here, we aimed to enhance the ecological validity of the learning situation: the ‘observer’ watched their friend (‘demonstrator’) performing a differential fear-conditioning task in real time. During the task, one conditioned stimulus (CS+) was repeatedly linked with electric stimulation (US) while another one (CS-) was always safe. Subsequently, the observer was presented with the CS+ and CS- directly but without receiving any shocks. Skin conductance (SCR) and fear-potentiated startle (FPS) responses were measured in observers throughout the whole experiment. While the US applied to the demonstrator elicited strong SCR in the observers, subsequent differential SCR to CSs (CS+ vs. CS-) presented directly were dependent on declarative knowledge of the CS+/US contingency. Contingency-aware observers also showed elevated FPS during both CS+ and CS- compared to intertrial intervals. We conclude that observational fear learning involves two components: an automatic emotional reaction to the response of the demonstrator and learning to predict stimulus contingency (CS+/US pairing). Ecological modifications proposed offer new perspectives on studying social learning of emotions.

## Introduction

Learning through interactions with others, that is, social learning, is often adaptive. For instance, receiving information about threats through social means helps to avoid costly first-hand experiences. Social transfer of information about threats is thought to be mediated by emotional contagion, a bottom-up process through which the emotional state of one organism elicits the same state in another ^1^. This basic ability to perceive and mimic the emotions of others is conserved across species, including apes ^2^, dogs ^3^ and birds ^4^. Further, the ability to not only imitate emotions but also learn from emotional expressions of others has been described in different species, such as rhesus monkeys ^5^ and rodents ^6,7^ In humans, it has been shown that watching an actor undergoing fear conditioning evokes vicarious reactions in the observer resulting in learning as measured by behavioral, psychophysiological, and neural responses ^8,9^ These responses can be modulated by a number of factors, including the observer’s empathy level ^10^, social group affiliation ^11^, and racial similarity to the actor ^12^.

In previous studies on vicarious fear learning in humans, participants observed an actor performing a differential fear conditioning task (in which one visual stimulus was repeatedly linked with aversive stimulation while another was always safe). Successful fear conditioning was typically assessed as an enhanced skin conductance response (SCR) to direct presentations of the stimulus which had been paired with an electric shock administered to the actor (further referred to as CS+). As most of these studies have used standardized video recordings of an anonymous person presented to individual participants, they have lacked ecological validity offered by dynamic interactions ^8,10^, but see ^13^.

In this study, we modified the experimental protocol proposed by Haaker and colleagues ^14^ to enhance its ecological validity and to test whether the modifications affected acquisition of fear. Ecologically valid (naturalistic) paradigms can be understood as involving realistic, interactive stimuli representative of real-world experiences while maintaining a reasonable degree of experimental control ^15,16^. We believe that modifying the procedure toward a more naturalistic one can tell us how observational fear learning occurs when authentic, individually varying emotions are being expressed. Consequently, for the purpose of adapting the protocol, we decided to invite pairs of participants and involve them both in the experimental procedure. Instead of using a prerecorded video of an actor, one of the two participants was asked to become a live demonstrator. Additionally, we decided to recruit pairs of friends expecting that it may increase learning efficiency, which has been shown to be enhanced when the learning model is perceived as similar, for example as he or she belongs to the same social group ^11,12^ Moreover, it has been shown that interpersonal liking increases emotional mimicry ^17^, which is yet another factor involved in emotional contagion ^18^. It has also been suggested that behaviors and emotional expressions of social ingroup members are mimicked preferentially, playing a role in social learning ^19^ Besides our motivation to examine the learning from authentically expressed emotions in friends in a dynamic social context, the changes proposed here are one step in narrowing the translational gap between the human observational fear conditioning paradigm and contemporary experiments on rodents, in which two interacting animals are typically used, e.g. ^6^. We hope that this can be used as a starting point for further studies comparing behavioral and neural correlates between rodent and human models of vicarious fear acquisition.

In our study, one of the participants (the demonstrator) was subjected to a differential fear conditioning task while being watched by another participant (the observer) through a live video stream. The skin conductance and fear-potentiated startle (FPS) responses of the observers were recorded as indicators of vicarious fear learning. Although both have been commonly used as measures of aversive conditioning, their underlying mechanisms differ ^20^. SCRs reflect sympathetic arousal, and they are elicited by salient or novel stimuli in general.

FPS relies on a simple defensive reflex, in which the primary reflex pathway is modulated by inputs from the amygdala ^21^, has been interpreted in terms of defensive reactivity. Unlike SCRs, FPS can be probed both during CS and between trials, the latter providing a baseline condition. Using FPS also provides a closer link to fear conditioning studies in rodents, where startle methodology is commonly applied and follows similar principles ^6^. Moreover, FPS has only recently been applied in the observational fear conditioning design ^22^ and no studies from this field have so far employed both these measures, making our study the first to incorporate both.

Our hypotheses were as follows: first, in the Observational Fear Learning (OFL) phase, we expected an augmented skin conductance response in the observers watching their friends receiving electric shocks. Second, we hypothesized that as a consequence of the pairing between CS+ and social US (friend’s reaction to the shock) watched during the learning phase, the observers would develop a conditioned response to the CS+ without directly experiencing the aversive stimulation. This socially driven conditioned reaction was expected to be reflected in stronger skin conductance and fear-potentiated startle responses after direct presentation of CS+ compared to CS-, when tested in the direct expression (DE) phase.

## Methods

### Participants

70 male volunteers (35 pairs of friends), aged between 18 and 27 (*M* = 21.4, *SD* = 2.2) participated in the study. Considering that primary observational fear conditioning effects reported previously, eg. ^11^ were large, our study was sufficiently powered to detect effects of similar size (see the Supplementary Methods for detailed explanation). To be eligible for the study, a pair had to have known each other for at least 3 years (in the recruited group: *M* =6.8, *SD* = 4.2) and score sufficiently high (minimum 30 out of 60 points; in the recruited group: *M* = 50.1, *SD* = 7.3) on the McGill Friendship Questionnaire ^23^. All the participants were screened for the ability to recognize the colors used in the task. Only heterosexual participants (based on self-declaration) were included. Within each pair, the subjects were randomly assigned roles – one person was the demonstrator (learning model) and the other one was the observer. The protocol of the study was approved by the Ethics Committee of the Faculty of Psychology at the University of Warsaw in accordance with the Declaration of Helsinki. All subjects gave their written informed consents and received financial compensation for participation in the study.

### Stimuli and Materials

Two large colored squares (blue and yellow) displayed initially on a computer screen in front of the learning model (demonstrator) and later in front of the observer (in OFL and DE stages respectively) served as conditioned stimuli (CS). The assignment of colors to CS+ (squares that might be reinforced with shocks) and CS- (squares that were never reinforced) was counterbalanced across participants. During intertrial intervals, a fixation cross was displayed. The OFL phase consisted of 24 CS+ and 24 CS-trials. During the course of the task, 12 CS+ were reinforced with an uncomfortable shock to the ventral part of the upper right forearm of the demonstrator (unconditioned stimulus, US). The CS+/CS- order was pseudo-random with the restriction that any given CS may not be repeated more than twice. The first and last presentation of the CS+ was always reinforced. Two sequences matching these criteria were created and counterbalanced across participants.

In each OFL trial, the CS was presented on the demonstrator’s screen for 9 seconds and the intertrial interval was randomized between 10 and 15 seconds. The US, administered in half of the CS+ presentations, started 7.5 seconds after CS onset and consisted of five unipolar electrical pulses delivered with 1 ms duration and a latency of 200 ms (total stimulation duration = 0.8 s). The resulting demonstrator’s reaction started straight after the stimulation administration and was likely to co-terminate with the end of the CS+ display. Electrode placement at the upper ventral part of the forearm was chosen because in most participants it caused muscle flexion and a resulting hand movement, even at non-painful stimulation intensities. The shock level was adjusted for each demonstrator individually prior to the learning session (see the Procedure), so that it was experienced as very uncomfortable but not painful. In order to measure the fear-potentiated startle response, the startle probes were presented to the observer during half of CSs (onsets, randomly chosen, at 6.0, 6.5 or 7.0 seconds after CS onset) and a quarter of intertrial intervals (onsets between 2.0 and 4.5 seconds after fixation cross onset, in order not to interfere with subsequent CS presentations). The acoustic startle probe was a white noise burst (80 dB(A) and 50 ms duration) presented binaurally through headphones. Although most studies investigating fear-potentiated reactivity have reported using louder stimuli (~90 dB), the volume that we used was high enough to elicit startle reflex ^24^ and at the same time not overly uncomfortable for the observers (which could interfere with the observational fear acquisition, especially given that the aversive unconditioned stimuli were never direct). The procedure, including number of trials, their timing and the reinforcement ratio were adjusted to match the demands of potential neuroimaging experiments and were based on existing recommendations ^14^.

The OFL phase was followed by the DE phase, which consisted of 12 CS+, 12 CS- (displayed on the observer’s screen), and no US. Here, timing of CSs, intertrial intervals, stimulus order and startle probe presentations followed the same rules as described above. A scheme presenting the experimental design is shown in Fig. 1.

**Figure 1.**
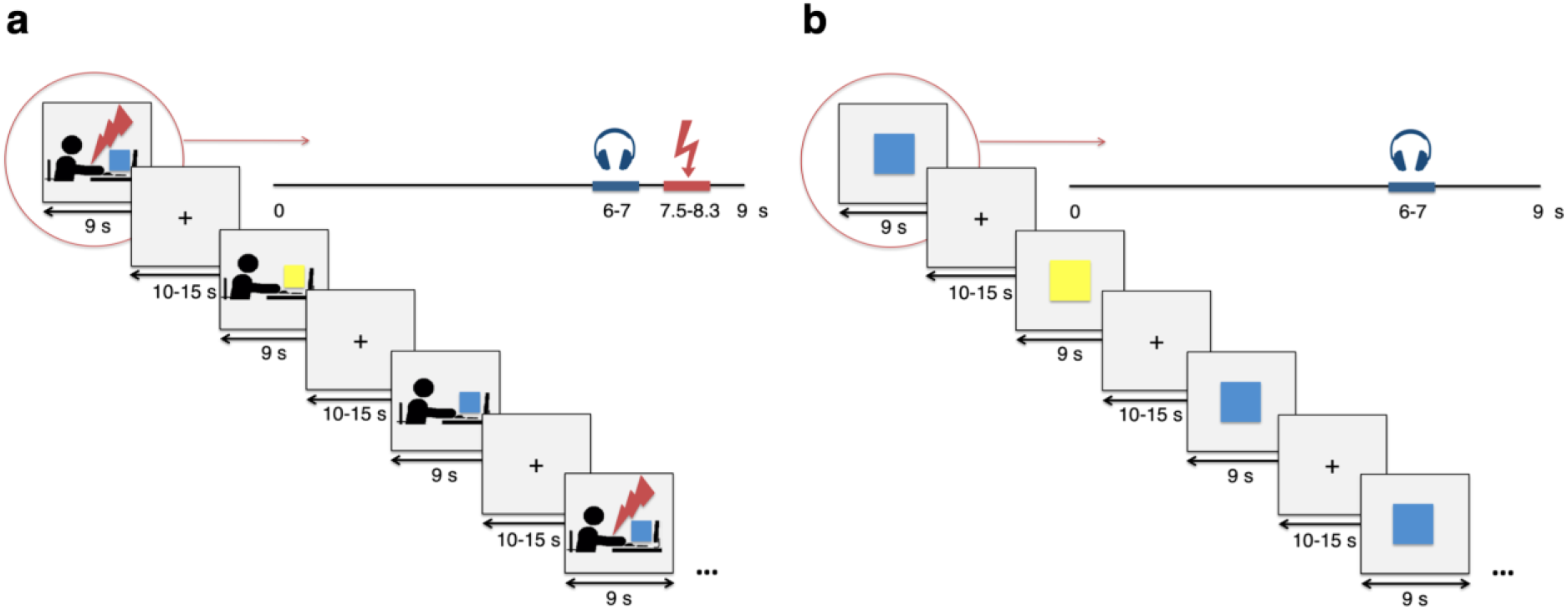
Design of the experiment. (**a)** In the OFL phase, the observer watched their friend performing a differential fear conditioning task. The conditioned stimuli (blue and yellow squares) were displayed on the demonstrator’s screen in a pseudorandom order. Presentation of one of the squares (CS+, here: blue) was accompanied with an uncomfortable electric shock (unconditioned stimulus, US) to the right forearm of the demonstrator with 50% probability. The other color square served as a stimulus that was never reinforced (CS-, here: yellow). A fixation cross was presented between the stimuli. The observer’s SCR and FPS response were used to assess learning. In order to measure the startle reflex magnitude, the white noise burst was pseudorandomly presented through the headphones in half of all types of trials (in which a blue square, yellow square, or fixation cross was presented). (**b)** In the DE phase, the observer performed the demonstrator’s task but the USs were not applied. Instructions provided to the observers did not suggest that the electrical stimulation would accompany only one CS in the OFL phase nor that it would be omitted in the DE phase.

For observation, an HD-SDI camera connected to a computer screen was used to transmit video (without sound) of one participant to the other. Stimulus presentation was controlled using Presentation v19.0 software (Neurobehavioral Systems, Berkeley, CA,

USA). For technical considerations regarding the equipment used, see the Supplementary Methods.

### Procedure

We used the experimental protocol of Haaker and colleagues ^14^ introducing the following modifications: both demonstrators and observers were invited to the laboratory to take part in the experiment, demonstrators were friends of the subjects, demonstration was transmitted via real time video streaming and fear-potentiated startle was included as an additional measure of conditioning. Upon arrival to the laboratory, the participants were shortly informed about the measurements and stimulation used in the experiment, they filled out the safety form to exclude contraindications for the electrical stimulation and signed the informed consent. Subsequently, they were randomly assigned to their roles as demonstrators and observers and invited to take a seat in one of two adjacent rooms. Skin conductance, electromyography (EMG, used for startle response measurement) and stimulation electrodes were attached and headphones were given to both participants. The demonstrator was seated slightly angled in relation to the computer screen, so that their face, stimulated hand and the computer screen were all visible to the camera relaying the image to the observer’s room. Next, both participants were informed about their roles (for detailed instructions see the Supplementary Methods). The demonstrator was told that they would perform a computer task involving colored figures and electric shocks and that their performance would be watched by their friend through the camera. Demonstrators were informed about the color of the figure that would be paired with electrical stimulation. Each demonstrator was also asked to signal each shock occurrence clearly and not try to stifle the discomfort reaction. Besides the natural twitch of the forearm, facial grimace was suggested as an accurate way of demonstrating discomfort. A pre-recorded video presenting an exemplary reaction was shown to each demonstrator in order to provide comparable expressions of discomfort across participants. At the same time, demonstrators were encouraged to behave as naturally as possible and to adjust their reactions so that they would be perceived as reliable. Providing such instructions enabled establishing a compromise between a fully spontaneous demonstrator’s behavior (which based on our qualitative observations from extensive pretests was not informative for the observers, as the reactions to non-painful pulses tended to be very subtle) and a fully controllable, carefully prepared videos. After the instructions, shock adjustment was performed. Stimulus intensity was gradually increased and after each trial the participant was asked to describe it using an 8-point scale, ranging from imperceptible (1) to painful (8). The adjustment was stopped when the participant described the shock as ‘very unpleasant, but not painful’ (6). At the time of the demonstrator’s preparations, the observer was seated in a separate room getting habituated to the noise bursts used as startle probes. Subsequently, the observer was informed about their role: they were asked to watch their friend performing a task involving colored figures and electric shocks and to ignore short loud sounds occurring occasionally. They were also told that after observation, they would do the same task themselves.

The OFL phase started with a repeat of the startle habituation (due to interference associated with instructions giving). When the OFL task was over, the video stream was turned off. The observer was then informed that it was their turn to do the same task and the DE phase followed. Identical stimulus material and parameters were used, however, no shocks were administered to the subjects. Finally, the observer filled out an online contingency questionnaire, with questions progressing from open-ended, through percentage ratings of the shock occurrence, to forced choice ^25^. Participants were classified as contingency-aware if their forced choice answer was correct and consistent with previous responses. Additionally, the observer was asked to rate the demonstrator’s performance in terms of the level of discomfort expressed as well as the strength and naturalness of reactions;also their level of empathy felt during the observation was assessed. Likert scales ranging from 0 (not at all; very poor) to 9 (very much; very strong) were used for these purposes. In the end, both participants were debriefed about the study.

### Physiological recordings, Data scoring and Analysis

Skin conductance and startle responses were collected only from the observers and sampled at 2 kHz (for details concerning the signal recording, see the Supplementary Methods).

#### Skin Conductance

The recorded signal was decomposed into tonic and phasic components using cvxEDA ^26^. Skin Conductance Responses were scored as the difference between the maximum value occurring between 0 and 6 seconds after stimulus onset and the mean value from the preceding 2 seconds. The response window was limited to 6 seconds in order to avoid entanglement of the CS and acoustic startle probe effects. While the phasic component should, by definition, have a zero baseline, the subtraction was done to avoid scoring spontaneous fluctuations occurring before the stimulus. Response window choice was a compromise between the recommendations of the original protocol (foot point of the responses at 0.5 to 4.5 s after stimulus ^14^), entire interval response approach ^27^ and the constraints imposed by startle application. Amplitude smaller than 0.2 μS was treated as no response. Due to timing proximity, responses to the US were measured only for trials containing no startle probes during and after CS presentation, and non-reinforced trials (no US) were also scored, by using an identical time window as when the stimulus was present.

SCR data from subjects for whom less than 5 non-zero responses were observed during the DE phase were excluded from the analysis (leading to exclusion of the SCR data of 3 subjects). The resulting amplitudes were normalized within subjects using a log-transformation: log (1 + SCR / SCR_max_), where SCR_max_ was the highest response found for a given subject. Mean magnitudes (including no-responses as zero amplitude) were calculated for each condition and each participant. Within-subject averages were submitted to group-level analysis.

#### Fear-Potentiated Startle

Following the guidelines ^24^, the signal was band-pass filtered in the 28 - 500 Hz band (4th order Butterworth filter), rectified (i.e. replaced with its absolute value), and smoothed using a 40 Hz low-pass FIR filter. Each response was scored as the difference between the maximum value in a 20 - 120 ms time window and mean value in the −100 - 0 ms time window (times relative to startle probe onset; trial was scored as 0 if peak amplitude was lower than baseline). Resulting magnitudes were normalized within participants using T-scores ^24^. Mean magnitudes were calculated for each participant and each condition and used for group-level analysis.

All trials were visually inspected for the presence of artifacts, which led to exclusion of data from one participant due to an overall noisy recording; however, no individual trials were discarded. EMG data from two other participants, exhibiting 5 or less non-zero responses throughout the entire experiment, were excluded from the analysis (note that different participants were excluded from the EMG and SCR analyses).

Custom Python scripts utilizing Numpy (https://www.numpy.org/), Scipy ^28^, and Bioread (github.com/uwmadison-chm/bioread) libraries were used for signal preprocessing and scoring of both skin conductance and startle EMG.

For the statistical analyses of psychophysiological responses, repeated measures and mixed-model ANOVAs as well as two-tailed paired t-tests were used. The main measures of interest were the conditioned responses (differences between reactions to CS+ and CS-, both skin conductance and startle) examined in the direct expression phase. PASW Statistics 18 (SPSS Inc., Chicago, IL) was used for statistical analyses.

## Results

### Behavioral results

Based on the criteria described above, participants’ contingency knowledge was assessed. A total of 14 out of 35 observers were classified as contingency-aware and 21 as contingency-unaware. The non-parametric Kruskal-Wallis test (*N* = 35) revealed no differences in the observers’ perception of the demonstrators between contingency-aware (*n* = 14) and -unaware (*n* = 21) groups regarding the 4 studied categories: discomfort expressed, strength of reaction, naturalness of reaction, and observer’s empathy (see Table 1).

**Table 1.**
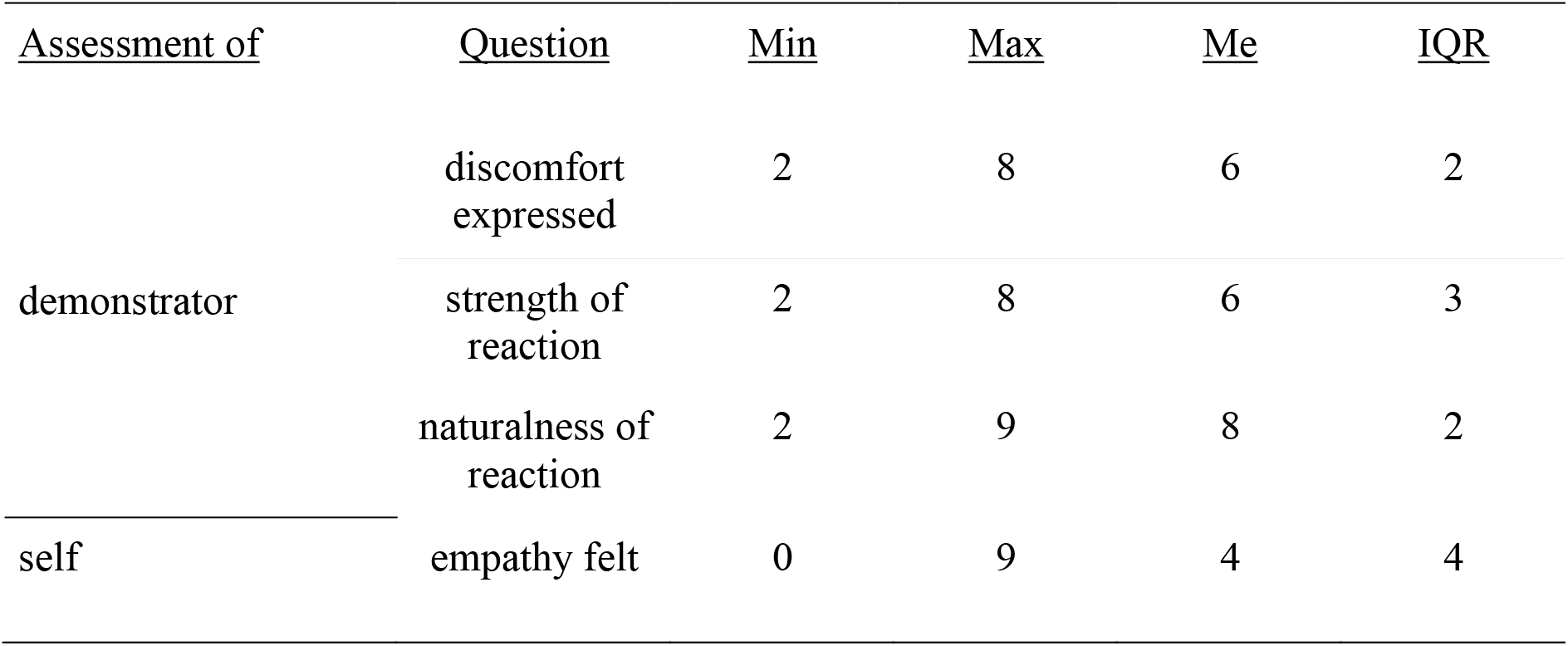
Observer’s post-experiment rating of the demonstrator’s expression as well as their self-empathy (felt during the observation phase) assessment. Me: median, IQR: interquartile range.

| <u>Assessment of</u> | <u>Question</u> | <u>Min</u> | <u>Max</u> | <u>Me</u> | <u>IQR</u> |
| --- | --- | --- | --- | --- | --- |
| demonstrator | discomfort expressed | 2 | 8 | 6 | 2 |
|  | strength of reaction | 2 | 8 | 6 | 3 |
|  | naturalness of reaction | 2 | 9 | 8 | 2 |
| self | empathy felt | 0 | 9 | 4 | 4 |

In the group of subjects eligible for SCR analysis, 14 of 32 observers were contingency-aware, while in the group selected for FPS analysis, 13 of 32 observers were contingency-aware.

### Skin conductance

In the whole group analysis (*N* = 32), a paired t-test revealed a significant difference between reactions measured when observers watched a friend receiving shocks and those recorded during observation of CS+ trials in which no shocks were applied, *t*(31) = 8.37, *p* < .001, *d* = 1.69 (see Fig. 2a). Regarding the SCRs to colored squares presented throughout the experiment, in a repeated-measures ANOVA (with stimulus type and task phase as within-subject factors) we observed no differential reactions to CS+ compared to CS-presentation either in the OFL or in the DE phase and no main effects were found.

**Figure 2.**
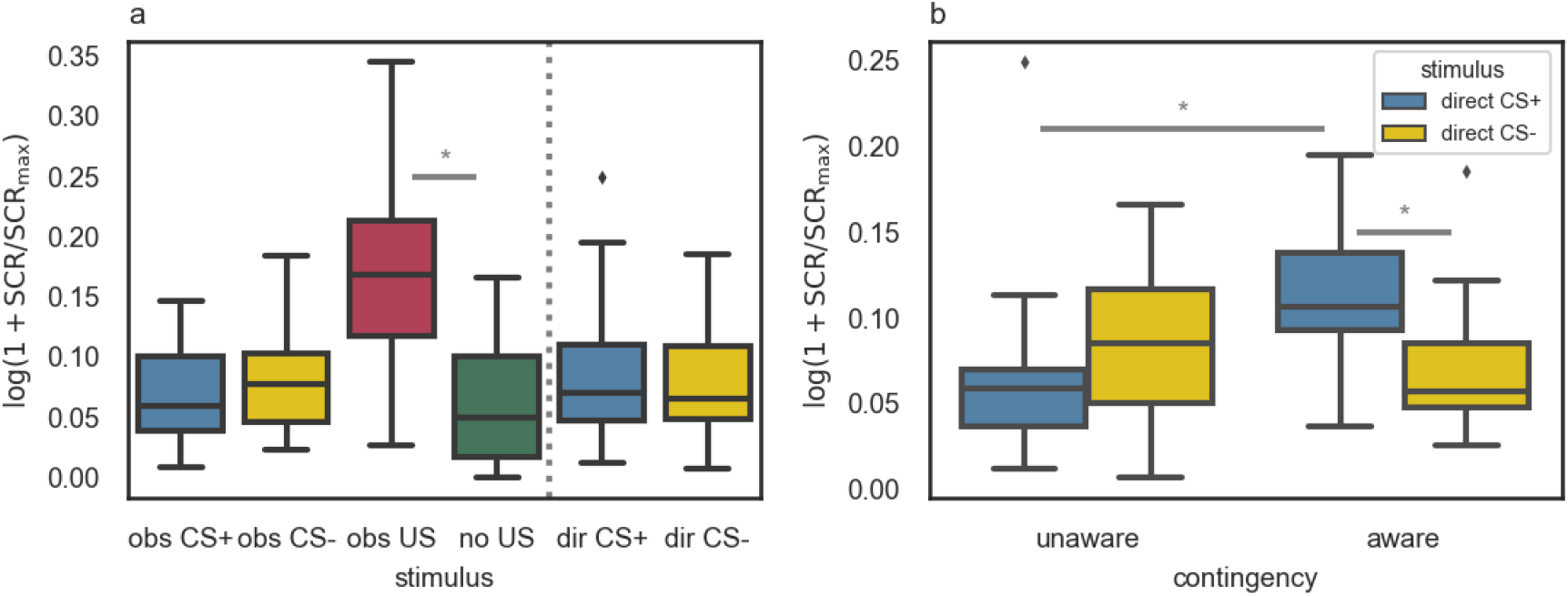
(**a**) SCR magnitudes to stimuli presented in the experiment: obs CS+ and obs CS- (appearance of CS+/CS- in the OFL phase), obs US and no US (observation of a friend receiving/not receiving electric shock during CS+ in the OFL phase), dir CS+ and dir CS- (appearance of CS+/CS- in the DE phase). Substantial differences are marked. *N* = 32. (**b**) SCR magnitudes to the appearance of the CS+ and CS- in the DE phase of the experiment for contingency-aware (*n* = 14) and -unaware (*n* = 18) participants separately. The stimulus (CS+ vs. CS-) x contingency (aware vs. unaware) interaction was found, *F*(1, 30) = 12.29, *p* = .001, η_p_^2^ = .29, and substantial effects of the post-hoc comparisons are marked. Error bars indicate 1.5*IQR beyond the 1^st^ quartile and above the 3^rd^ quartile. The * symbol was used for *p* < 0.05.

Having obtained a low ratio of the CS+/US contingency, we decided to check whether the contingency knowledge moderates the relationship between SCRs and the conditioned stimuli (CS+ and CS-). For this purpose, we used a mixed model ANOVA with the skin conductance response as a dependent variable, stimulus type as a within-subject factor, and contingency knowledge as a between-subject factor. In the direct expression phase we found the stimulus (CS+ vs. CS-) x contingency knowledge (known vs. unknown) interaction, *F*(1, 30) = 12.29, *p* = .001, η_p_^2^ =.29. Post hoc comparisons (with Bonferroni correction) revealed a stronger skin conductance response to CS+ (*M* = 0.11, *SD* = 0.05) as compared to CS- (*M* = .07, *SD* = 0.04, *p* = .002, *d* = 0.88, 95% CI [0.02, 0.07]) in participants who indicated the CS+/US contingency correctly. Moreover, skin conductance response to CS+ was stronger in the group of contingency-aware (*M* = 0.11, *SD* = 0.05) compared to -unaware participants (*M* = 0.07, *SD* = 0.05, *p* = .015, *d* = 0.80, 95% CI [0.01, 0.08]; see Fig. 2b).

### Fear-potentiated startle

In the whole group analysis (*N* = 32), in order to check for the differences in FPS responses to different stimuli depending on the phase of the experiment, we ran a repeated measures ANOVA with the startle response as a dependent variable and stimulus type and task phase as within-subject factors. We found the main effect of phase, *F*(1, 31) = 81.87, *p* < .001, η_p_^2^ = .73. Post hoc comparisons (with Bonferroni correction) revealed a stronger FPS response in the DE (*M* = 54.14, *SD* = 2.59) compared to the OFL phase (*M* = 47.93, *SD* = 1.30,*p* < .001, *d* = 3.03, 95% CI [4.81, 7.62]; see Fig. 3a).

**Figure 3.**
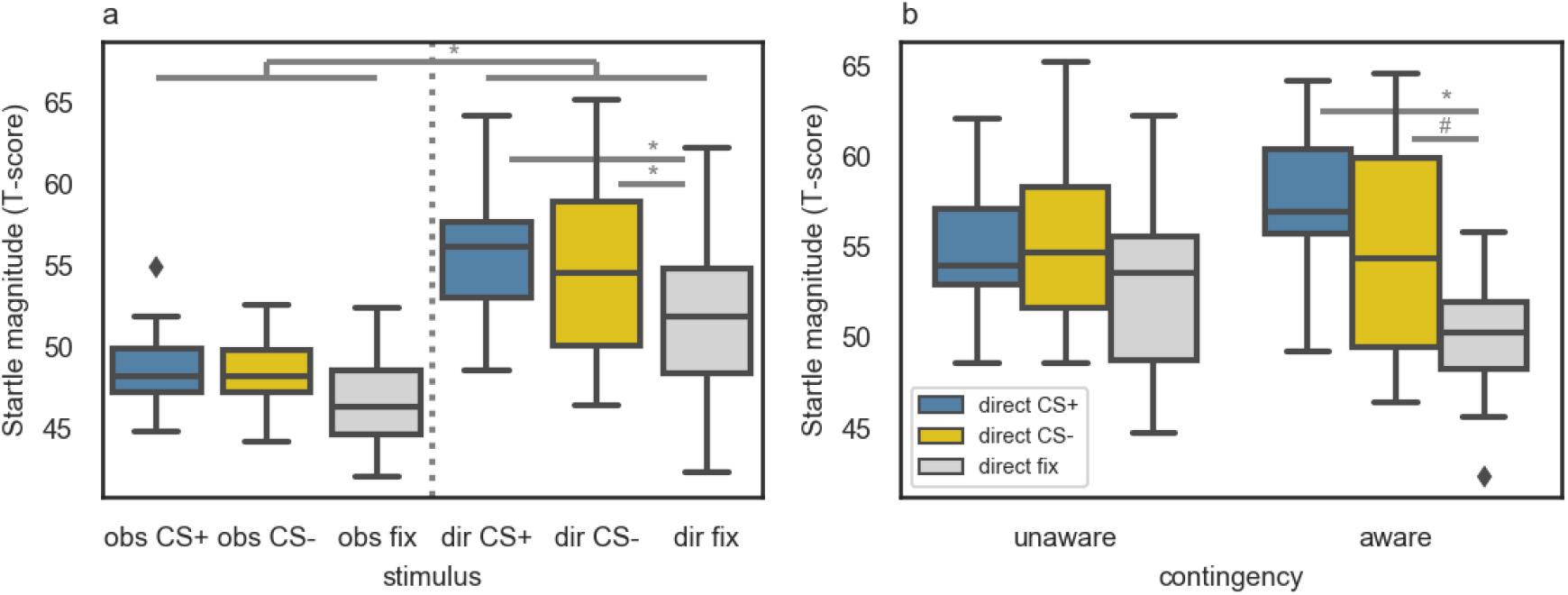
(**a**) FPS responses measured during different stimuli presented in the experiment: obs CS+, obs CS- and obs fix (presentation of CS+/CS-/fixation cross in the OFL phase), dir CS+, dir CS- and dir fix (presentation of CS+/CS-/fixation cross in the DE phase). Substantial differences are marked. *N* = 32. (**b**) FPS magnitudes measured in the DE phase: results of contingency-aware (*n* = 13) and -unaware (*n* = 19) participants are grouped separately. A trend toward the stimulus (CS+ vs. CS-vs. fixation cross) x contingency (aware vs. unaware) interaction was found, *F*(2, 60) = 2.59, *p* = .083, η_p_^2^ = .08, and substantial effects of the post-hoc comparisons are marked. Error bars indicate 1.5*IQR beyond the 1^st^ quartile and above the 3^rd^ quartile. The * symbol was used for*p* < 0.05, and # symbol for *p* < 0.1.

To check whether there was an effect of contingency knowledge on FPS responses to different stimuli in the DE phase, we performed a mixed model ANOVA analogous to the analysis of the SCR data. We found a significant main effect of stimulus (CS+ vs. CS-vs. fixations cross), *F*(2, 60) = 8.16, *p* = .001, η_p_^2^ = .21. Further post hoc comparisons (with Bonferroni correction) showed that the FPS reaction to both CS+ (*M* = 55.71, *SD* = 4.08) and CS- (*M* = 54.99, *SD* = 5.21) was stronger than the one measured during the fixation cross presentation (*M* = 51.73, *SD* = 4.55); for CS+ > fixation cross: *p* < .001, *d* = 0.92, 95% CI [1.97, 6.99]; for CS- > fixation cross: *p* = .034, *d* = 0.67, 95% CI [0.22, 6.85]; see Fig. 3A. There was also a trend toward the stimulus (CS+ vs. CS- vs. fixation cross) x contingency knowledge (aware vs. unaware) interaction, *F*(2, 60) = 2.59, *p* = .083, η_p_^2^ = .08. Results of the post hoc comparisons (with Bonferroni correction) revealed that in the group of contingency-aware participants, the response to both CS+ (*M* = 57.13, *SD* = 4.56) and CS- (*M* = 55.01, *SD* = 6.17) was stronger than the response to a fixation cross (*M* = 49.10, *SD* = 3.85); for CS+ > fixation cross: *p* < .001, *d* = 1.69, 95% CI [3.27, 11.00]; for CS- > fixation cross: *p* = .056, *d* = 0.97, 95% CI [-0.10, 10.12]; see Fig. 3b. This trend suggests that the effect of the potentiated startle response to both conditioned stimuli might be specific to only the contingency-aware participants.

## Discussion

In the present study the observational fear conditioning was tested in a paradigm with improved ecological validity. Most previous studies on this phenomenon have used prepared video recordings with actors unknown to the observers. Instead, we arranged a real-time experiment involving participants who were significant to each other. Further, to better reflect activity of the defensive system, in addition to SCR, we used FPS. As the latter is commonly used in rodent studies, it additionally allows for across species comparisons. Both measures confirmed that acquisition of fear through observation was successful. They also showed that differential fear learning was dependent on declarative knowledge of the CS+/US contingency.

We were particularly interested in the results obtained during the direct expression (DE) phase, as they reflected the observational fear learning efficiency. These results confirmed that the observers learned to respond with physiologically stronger reactions to threat, but only when they were aware of the CS-US contingency. Specifically, the contingency-aware participants responded with stronger SCRs to the direct CS+ (vs. CS-) and increased startle potentiation towards both CSs (vs. fixation cross). The latter effect suggests that the aversive learning generalized; indeed, it is possible that during the OFL phase participants learned to associate aversive feelings with both squares, as opposed to the fixation cross, and, as a consequence, during the DE phase, both CSs (and not the fixation cross) elicited augmented startle responses.

When taking a closer look at the SCR results, two effects are apparent. First, observational US elicited stronger and more robust reactions than any of the conditioned stimuli in the DE phase. This means that the US was perceived the way we expected – which is important, considering that observing the aversive qualities of the US is a prerequisite for threat learning. Second, differential conditioning of the SCRs was observed only in the contingency-aware participants. This observation could be made because the low rate of contingency awareness declared in the post-experimental questionnaire allowed splitting the participants into two groups and it suggests that learning about CS-US contingency can be separated from the response to the observational US.

In the case of FPS results, we observe several effects. Overall, the reactions were stronger in the DE compared to the OFL stage, suggesting that the an elevated sense of threat was induced during this part of the experiment, and that following observation the observers indeed expected to experience the aversive stimulation themselves. This remains in line with the interpretation of startle response as reflecting defensive preparation or reactivity to stimuli suggesting direct threat. Furthermore, in the DE stage, startle probes presented during both conditioned stimuli elicited heightened responses compared to the ones measured during intertrial intervals, suggesting that the acquired fear was generalized (nonspecific). We observed a trend-level interaction with contingency knowledge, which upon inspection suggests that the effect is driven by the contingency-aware participants. Although the SCR and FPS results did not show one-to-one correspondence, the information they provide is complementary and consistent. It seems that both attention and defensive reactivity are increased as a consequence of observational fear conditioning in the tested paradigm and that these effects are specific for participants reporting declarative knowledge about the CS+/US relationship.

The physiological results replicate the findings of previous works which manipulated various elements in a video-based vicarious learning procedure and consistently showed differential conditioning effects measured by SCR ^10–12^ and FPS ^22^ Interestingly, in the aforementioned studies, an overwhelming majority of the participants correctly identified the CS+/US contingency and the few who did not were excluded from analyses ^29^. In our experiment, the general learning efficiency (declarative contingency knowledge) was low, with 14 out of 35 participants correctly identifying the relationship between CS and US. Therefore, the largest difference observed in the current results lies in the low ratio of the CS+/US contingency observed.

The changes we introduced to the reference protocol ^14^ were designed to increase its ecological validity. To improve the immediacy and realism of the threat learning situation, we invited both demonstrator and observer (who were well known to each other) to the laboratory and carried out the observational stage via live streaming. At the same time, the increase of realism came at a cost of lessening our control over experimental conditions: although the demonstrators received general instructions about how they should react to the unpleasant stimulus, each of them still reacted in their own individual way. Such reactions could not be influenced once the experiment started and were not selected through video editing before the experiment ^14^ While this improved realism, it might have also increased ambiguity (leading to lowered rate of contingency awareness), since spontaneous displays of emotions are smaller in amplitude and have different internal timing than posed emotional expressions ^17^ The ambiguity, however, should have been reduced by the fact that the observers were familiar with the demonstrators and thus more likely to be sensitive to their emotional expressions. While our ecologically-aimed modifications likely contributed to the low contingency knowledge, other factors might come into play as well. First, the inclusion of startle probes ^30^ and low (50%) CS+ reinforcement ratio might have further contributed to the creation of a weak learning situation. It is possible that auditory stimuli not only disrupted participants’ attention but also introduced an additional aversive component which interfered with the clear CS+/US association. Increase of the CS+ reinforcement ratio should be considered as a possibility for the contingency learning enhancement. Furthermore, the demonstrators were seated in an ordinary room, rather than against a clear and contrasting background, which might have introduced additional visual distractors.

We find the low level of contingency knowledge and its impact on conditioning measures interesting (as it highlights the role of the observer’s attention and the observational stimulus quality), but not surprising, given that the role of contingency knowledge has been described for various conditioning protocols and conditioning measures, e.g. ^25,31,32^ Skin conductance response has been reported as a measure reflecting contingency learning ^33^, while potentiation of the acoustic startle reflex has been claimed to be more valence-specific and less dependent on attention ^34^, although see ^35^, but its relation to contingency awareness is still unclear. We showed that under ecological conditions, contingency learning is reflected in augmented physiological reactions to potential threat, as indicated by stronger skin conductance and fear potentiated startle responses of the contingency-aware observers. Thanks to employing two different psychophysiological measures of conditioning, we demonstrated two effects: differentiation of the SCR and generalization of the FPS response, both relying on contingency knowledge. Our findings indicate an important characteristic of the human defensive system, suggesting its high sensitivity to all the cues related to the potential threat as well as its dependence on conscious processing of the context.

It is important to emphasize the significance of this study for further development of the vicarious fear learning research. The design proposed here is conceptually similar to the direct threat models of observational fear conditioning in rodents ^36,37^. The main advantage of our design is the real involvement of both demonstrator and observer as well as close relationship between both participants (in rodent studies, subjects are typically cage mates). Furthermore, acoustic startle response is commonly measured in rodent studies and its underlying neural circuit, involving the brainstem and centromedial amygdala, is conserved across species ^21,38^. These commonalities give rise to a whole range of possibilities for studying mechanisms of social fear acquisition across species. Research questions that might be answered owing to translational studies involve, for example, comparison of neural circuits related to observational fear learning in humans and rodents, investigation of the impact of participants’ familiarity on social fear learning (on both behavioral and neural level), and examination of different channels used for fear transmission (e.g. odor communication in humans).

One of the main limitations of the current study is a relatively small sample size, especially when group comparisons are considered. Although the previous reports suggested that the primary effects are strong and that reliable results should be observed also for current size subgroups, the interaction effects we observed are small. Nevertheless, our results are consistent with previous findings and validate the methodological frame for further ecological studies.

Arguably, the interaction between two participants could have been more natural if they were in the same room throughout the experiment. However, considering that the participants started the experiment together, we believe that their sense of involvement was maintained and likely much higher than in the case of recorded actors. At the same time, video streaming is more viable for different applications, such as fMRI or eye tracking. Thus, the proposed design constitutes a trade-off between naturalistic conditions and methodological requirements of potential applications. Finally, one aspect which the current study left unaddressed was the effects of specific dyads, for example due to relationship strength ^39–41^, on observational learning. We believe it might be addressed more accurately by further research, with larger and more varied groups of participants.

To conclude, in this work we show that participants watching their friends acquiring fear in real time can learn about the observed threat vicariously although the learning efficiency is lower than reported in previous studies. On the other hand, we found that the defensive reactivity of all the observers increased following the observational fear learning phase and this emotional reaction was generalized to both conditioned stimuli. The results suggest that the process of learning fear through observation may be composed of two separable components: an automatic, non-specific emotional reaction to the response of the demonstrator (which can serve as social unconditioned stimulus, US) and learning about predicting stimulus (CS) - US contingency. Further studies are needed to describe the factors underlying successful observational fear learning under ecological conditions.

## Data availability

Data from psychophysiological recordings are stored in an OSF repository and are available at https://osf.io/d3wxn/?view_only=ad87fa2e0076487c92c905f3be933720 [note: this is a view only link for review, OSF project will be made public upon publication]. Code replicating analyses reported here is available at https://github.com/mslw/vic-fear-learning-physio.

## Acknowledgements

We would like to thank Artur Marchewka for his valuable comments regarding the study design and the interpretation of results. Data collection and analysis were sponsored by National Science Centre grant 2015/19/B/HS6/02209. Ewelina Knapska was supported by European Research Council Starting Grant (H 415148).

## Author contributions statement

All the authors participated in designing the experiment, M. S. and A. K. conducted the experiment, performed the data analysis and drafted the manuscript, and J. M. M., M. W., A.O. and E. K. provided revisions. All authors approved the final version of the manuscript for submission.

## Additional information

The authors declare no competing interests.

