## Supplementary materials for "Observed but Never Experienced – Vicarious Learning of Fear Under Ecological Conditions"

### Methods

#### *Instructions given to the participants (translation)*

The demonstrator's role explanation: *You are the demonstrator. It means that you will be asked to do the task and your performance will be observed by your friend. In the task you will be shown colored figures but only one of them – the blue/yellow one* (depending on the experiment version) *– will be paired with an electric shock administered to the forearm in 50 percent of appearances. It will be very unpleasant but not painful. Please signal the shock occurrences clearly, so that your friend knows that they cause your discomfort. Besides the natural twitch of the forearm, please also try to react with a facial grimace of discomfort, e.g. squint your eyes and frown your forehead. It is very important that you react distinctly only in response to the shock administration – please try not to react to any other stimuli.*

The observer's role explanation: *You are the observer. It means that you will be asked to watch your friend performing the task. During the task both of you will see colored figures presented on his computer screen and hear short, loud sounds. The sounds will not be associated with the task though, so please try to ignore them. Your friend will receive electric shocks several times during the task. The shocks will be very unpleasant but not painful and they will induce certain reaction in your friend. Please focus on your friend performing the task. Later on, you will be asked to do the same task yourself.*

#### *Note on sample size*

In a recent SCR study examining effects of social group biases on observational fear conditioning 1, the effect size in the related samples t-test (CS+ vs CS- during the early stage of the direct test) reported for the subgroup of observers most comparable with the current study (racial and social in-groups with the demonstrator) and calculated based on supplementary data was d = 0.82. By convention, such effect is considered as large. Assuming d = 0.82, a sample size of just 14 subjects is sufficient to achieve 80 % power, as calculated using G*Power.

#### *Technical considerations*

Electrical stimulation was delivered using Biopac STM100C and STMISOC stimulator modules, driven by a National Instruments USB-6001 analog output card. Two Ag/AgCl electrodes placed 3.5 cm apart (measured between centers) and filled with salt free electrode gel were used for that purpose.

Using an HD-SDI security camera with an HDMI converter has an advantage of low latency and high robustness: SDI signal can be transmitted over long distances using a concentric cable, such setup is easy to connect and does not involve a PC for video processing. Using an action camera with direct HDMI output can be equally convenient, but unamplified HDMI cables are usually limited to 15 meters, which should be taken into consideration. In both cases, demonstrator’s screen brightness and camera white balance should be carefully adjusted to ensure accurate representation of conditioned stimuli.

#### *Physiological recordings*

Electrophysiological measurements were collected only from the observers. A Biopac MP-160 acquisition system and AcqKnowledge 4.0 software (Biopac Systems Inc., Goleta, CA, USA) were used to collect the data. Skin conductance was recorded using a Biopac EDA100C amplifier (gain: 5µS/V, 10 Hz low-pass hardware filter) and sampled at 2 kHz. Two 6 mm Ag-AgCl electrodes were attached to the distal phalanges of the second and third digits of the participant's left hand. Startle response was recorded using a Biopac EMG100C amplifier (gain: 2000, 1 Hz high-pass, 500 Hz low-pass hardware filter) and sampled at 2 kHz. Two Ag-AgCl shielded electrodes were attached under participant's right eye, on the orbicularis oculi muscle (one electrode in line with the pupil in forward gaze, the second 1-2 cm laterally). The ground electrode (identical, except non-shielded) was placed on the participant's forehead, directly under the hairline.
